# Dual Competition between the Basal Ganglia and the Cortex: from Action-Outcome to Stimulus-Response

**DOI:** 10.1101/187294

**Authors:** Meropi Topalidou, Daisuke Kase, Thomas Boraud, Nicolas P. Rougier

**Affiliations:** INRIA Bordeaux Sud-Ouest 33405 Talence, France; Institut des Maladies Neurodégénératives, Université de Bordeaux, 33000, Bordeaux, France; Institut des Maladies Neurodégénératives, CNRS, UMR 5293, 33000, Bordeaux, France; LaBRI, Université de Bordeaux, Institut Polytechnique de Bordeaux, CNRS, UMR 5800, 33405 Talence, France; CNRS, French-Israeli Neuroscience Lab, 33000 Bordeaux, France; CHU de Bordeaux, Institut MN Clinique, 33000 Bordeaux, France

**Keywords:** Cortex, Basal Ganglia, Competition, Short-range Excitation, Long-range Inhibition, Segregated Loops, Direct Pathway, Hyperdirect Pathway, Reinforcement Learning, Hebbian Learning, Covert Learning, Transfer Learning, Stimulus-Response, Action-Outcome

## Abstract

Action-outcome (A-O) and stimulus-response (S-R) processes that are two forms of instrumental conditioning that are important components of decision making and action selection. The former adapts its response according to the outcome while the latter is insensitive to the outcome. An unsolved question is how these two processes emerge, cooperate and interact inside the brain in order to issue a unique behavioral answer. Here we propose a model of the interaction between the cortex, the basal ganglia and the thalamus based on a dual competition. We hypothesize that the striatum, the subthalamic nucleus, the internal pallidum (GPi), the thalamus, and the cortex are involved in closed feedback loops through the hyperdirect and direct pathways. These loops support a competition process that results in the ability for the basal ganglia to make a cognitive decision followed by a motor decision. Considering lateral cortical interactions (short range excitation, long range inhibition), another competition takes place inside the cortex allowing this latter to make a cognitive and a motor decision. We show how this dual competition endows the model with two regimes. One is oriented towards action-outcome and is driven by reinforcement learning, the other is oriented towards stimulus-response and is driven by Hebbian learning. The final decision is made according to a combination of these two mechanisms with a gradual transfer from the former to the latter. We confirmed these theoretical results on primates using a two-armed bandit task and a reversible bilateral inactivation of the internal part of the globus pallidus.

## Introduction

Action-outcome (A-O) and stimulus-response (S-R) processes that are two forms of instrumental conditioning and important components of behavior. The former evaluates the benefit of an action in order to choose the best action among those available (action selection) while the latter is responsible for automatic behavior (routines), eliciting a response as soon as a known stimulus is present (Mishkin, Malamut, & Bachevalier, 1984; Graybiel, 2008), independently of the hedonic value of the stimulus. Action selection can be easily characterized using a simple operant conditioning setup such as for example, a two-armed bandit task where an animal must choose between two options of different value, the value being probability, magnitude or quality of reward (Pasquereau et al., 2007; Guthrie, Leblois, Garenne, & Boraud, 2013). After some trials and errors, a wide variety of vertebrates are able to select the best option (Herrnstein, 1974; Graft, Lea, & Whitworth, 1977; Matthews & Temple, 1979; Bradshaw, Szabadi, Bevan, & Ruddle, 1979; Dougan, McSweeney, & Farmer, 1985; Herrnstein, Vaughan, Mumford, & Kosslyn, 1989; Lau & Glimcher, 2005, 2008; Gilbert-Norton, Shahan, & Shivik, 2009). This selection is believed to result from the behavioral expression of the action-selection system. If the associated values are to be changed after only a few trials, the animal can still adapt its behavior and select rapidly the new best option. However, after intensive training (that depends on the species and the task) and if the same values are used all along, the animal will tend to become insensitive to change and persist in selecting the formerly best option (Lau & Glimcher, 2005; Yin & Knowlton, 2006).

Most of the studies on action selection and habits/routines agree on a slow and incremental transfer from the action-outcome to the stimulus-response system such that after extensive training, the S-R system takes control of behavior and the animal becomes insensitive to reward devaluation (Packard & Knowlton, 2002; Seger & Spiering, 2011). But very little is known on the exact mechanism underlying such transfer and one difficult question that immediately arises is when and how the brain switches from a flexible action selection system to a more static one. Our working hypothesis is that there is no need for such an explicit switch. We propose instead that an action expressed in the motor area results from both the continuous cooperation (acquisition) and competition (expression) of the two systems.

To do so, we consider the now classical actor-critic model of decision making elaborated in the 1980s that posits there are two separate components in order to explicitly represent the policy independently of the value function. The actor is in charge of choosing an action in a given state (policy) while the critic is in charge of evaluating (criticizing) the current state (value function). This classical view has been used extensively for modelling the basal ganglia (Suri, R E & Schultz, W, 1999; Suri, 2002; Frank, 2004; Doya, 2007; Glimcher, 2011; Doll, Bradley B, Simon, Dylan A, & Daw, Nathaniel D, 2012) even though the precise anatomical mapping of these two components is still subject to debate and may diverge from one model to the other (Redgrave, Peter, Gurney, Kevin, & Reynolds, John, 2008; Niv, Yael & Langdon, Angela, 2016). However, all these models share the implicit assumption that the actor and the critic are acting in concert, i.e. the actor determines the policy exclusively from the values estimated by the critic, as in Q-Learning or SARSA. Interestingly enough, (Sutton, R S & Barto, A G, 1998) noted in their seminal work that one could imagine intermediate architectures in which both an action-value function and an independent policy would be learned. We support this latter hypothesis based on a decision-making model that is grounded on anatomical and physiological data and that identify the cortex-basal ganglia (CBG) loop as the actor. The critic — of which the Substantia Nigra pars compacta (SNc) and the Ventral Tegmental Area (VTA) are essential components — interacts through dopamine projections to the striatum (Leblois, Boraud, Meissner, Bergman, & Hansel, 2006). Decision is generated by symmetry breaking mechanism that emerges from competitions processes between positives and negatives feedback loop encompassing the full CBG network (Guthrie et al., 2013). This model captured faithfully behavioural, electrophysiological and pharmacological data we obtained in primates using implicit variant of two-armed bandit tasks — that assessed both learning and decision making — but was less consistent with the explicit version (i.e. when values are known from the beginning of the task) that focus on the decision process only.

We therefore modified this early model by adding a cortical module that has been granted with a competition mechanism and Hebbian learning (Doya, 2000). This improved version of the model predicts that the whole CBG loop is actually necessary for the implicit version of the task, however, when the basal ganglia feedback to cortex is disconnected, the system is still able to choose in the explicit version of the task. Our experimental data fully confirmed this prediction (Piron et al., 2016) and allowed to solve an old conundrum concerning the pathophysiology of the BG which was that lesion or jamming of the output of the BG improve Parkinson patient motor symptoms while it affects marginally their cognitive and psycho-motor performances. An interesting prediction of this generalized actor-critic architecture is that the valuation of options and the behavioural outcome are segregated. In the computational model, it implies that if we block the output of the basal ganglia in a two-armed bandit task before learning, and because reinforcement learning occurred at the striatal level under dopaminergic control, this should induce covert learning when the model chooses randomly. The goal of this study is thus twofold: i) to present a comprehensive description of the model in order to provide the framework for an experimental paradigm that allow to unravel covert learning and ii) to test this prediction in monkeys.

## Materials and Methods

### The task

We consider a variant of a n-armed bandit task (Katehakis & Veinott, 1987; Auer, Cesa-Bianchi, Freund, & Schapire, 2002) where a player must decide which arm of *n* slot machines to play in a finite sequence of trials such as to maximize his accumulated reward. This task has received much attention in the literature (e.g. machine learning, psychology, biology, game theory, economics, neuroscience, etc.) because it provides a simple model to explore the trade-off between exploration (trying out a new arm to collect information about its payoff) and exploitation (playing the arm with the highest expected payoff)(Robbins, 1952; Gittins, 1979). This task has been shown to be solvable for a large number of different living beings, with (Plowright & Shettleworth, 1990; Keasar, 2002; Steyvers, Lee, & Wagenmakers, 2009) or without a brain (Reid et al., 2016), and even a clever physical apparatus can solve the task (Naruse et al., 2015).

#### The computational task

In the present study, we restrict the n-armed bandit task to *n* = 2 with an explicit dissociation between the choice of the option (*cognitive* choice) and the actual triggering of the option (*motor* choice). This introduces a supplementary difficulty because only the motor choice – the physical (and visible) expression of the choice – will be taken into account when computing the reward. If cognitive and motor choices are incongruent, only the motor choices matters. Unless specified otherwise, we consider a set of cues {*C*_*i*_}_i∈[1,*n*]_ associated with reward probabilities {*P*_*i*_}_i∈[1,*n*]_ and a set of four different locations ({*L*_*i*_}_i∈[1,4]_) corresponding to the *up*, *down*, *left*, *right* positions on the screen. A trial is made of the presentation of two random cues *C*_*i*_ and *C*_*j*_ (*i* ≠ *j*) at two random locations (*L*_*i*_ and *L*_*j*_) such that we have *L*_*i*_ ≠ *L*_*j*_ (see Fig. 1). A session is made of *n* successive trials and can use one to several different cue sets depending on the condition studied (e.g. reversal, devaluation). Unless specified otherwise, in the present study, exactly one cue set is used throughout a whole session.

**Figure 1.**
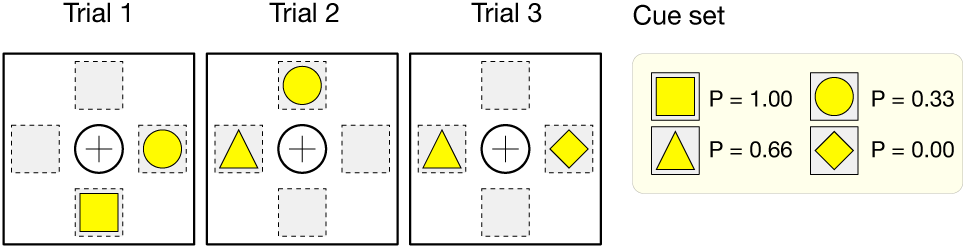
Three task trials from a 4-items cue set (◻, △, ◯, ◊) with respective reward probabilities (1,⅔,⅓, 0).

Once a legal motor decision has been made, reward is computed by drawing a random uniform number between 0 and 1. If the number is less or equal to the reward probability of the chosen cue, a reward of 1 is given, else, a reward of 0 is given. If no motor choice has been made or if the motor choice leads to an empty location (illegal choice), the trial is considered to be failed and no reward is given, which is different from giving a reward of 0. Best choice for a trial is defined as the choice of the cue associated with the highest reward probability among the two presented cues. Performance is defined as the ratio of best choices over the total number of trials. A perfect player with full-knowledge can achieve a performance of 1 while the mean expectation of reward is directly dependent on the cue sampling policy^1^.

#### The behavioral task

*With kind permission from the authors (Piron et al., 2016), we reproduce here the details of the experimental task which is similar*.

The primates were trained daily in the experimental room and familiarized with the setup, which consisted of 4 buttons placed on a board at different locations (0°, 90°, 180°, and 270°) and a further button in a central position, which detects contact with a monkey’s hand. These buttons correspond to the 4 possible display positions of a cursor on a vertical screen. The monkeys were seated in chairs in front of this screen at a distance of 50cm (Fig. 2). The monkeys initiated a trial by keeping their hands on the central button, which induced the appearance of the cursor in the central position of the screen. After a random delay (0.5s to 1.5s), 2 cues appeared in 2 (of 4) different positions determined randomly for each trial. Each cue had a fixed probability of reward (*P*_1_ =0.75 and *P*_2_ = 0.25) and remains the same same during a session. Once the cues were shown, the monkeys had a random duration time window (0.5s to 1.5s) to press the button associated with one cue. It moves the cursor over the chosen cue and they have to maintain the position for 0.5 s to 1.5 s. After this delay, the monkeys were rewarded (0.3 ml of water) or not according to the reward probability of the chosen target. An end-of-trial signal corresponding to the disappearance of the cursor was given, indicating to the monkeys that the trial was finished and they could start a new trial after an inter-trial interval between 0.5 s and 1.5s.

**Figure 2.**
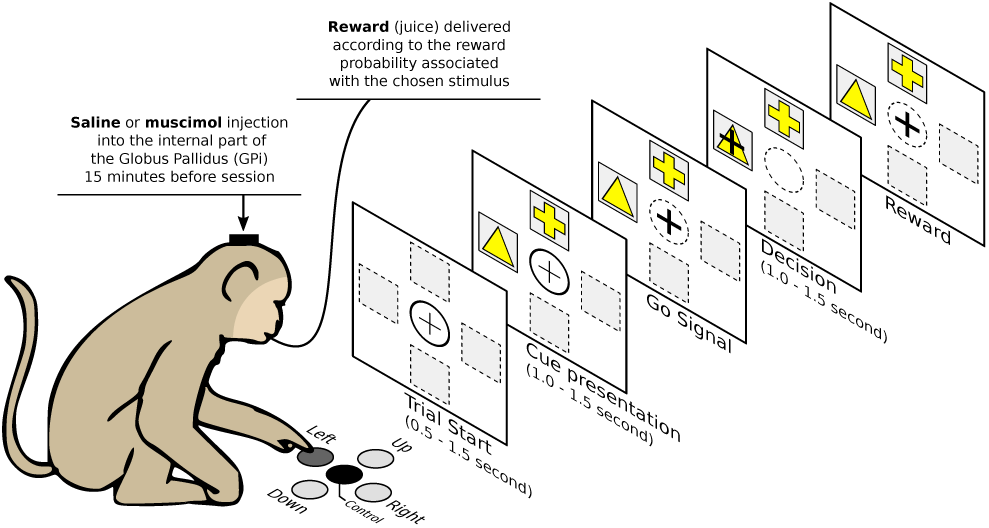
The behavioral task. The monkeys initiate a trial by keeping their hands on the central button, which induced the appearance of the cursor in the central position of the screen. After a random delay, two cues appears in 2 different positions. The monkey has a random duration time window (0.5s to 1.5s) to press the button associated with one cue. It moves the cursor over the chosen cue and it has to maintain the position for some duration. After this delay, the monkey is rewarded (0.3 ml of water) or not according to the reward probability of the chosen cue.

### The model

The model is designed to study the implications of a dual competition between the cortex and the basal ganglia (BG). The competition inside the cortex is conveyed through direct lateral interactions (short-range excitation and long range inhibition, (H. R. Wilson & Cowan, 1972, 1973; Coultrip, Granger, & Lynch, 1992; Muir & Cook, 2014; Deco et al., 2014)) while the competition within the BG is conveyed through the direct and hyperdirect pathways (Leblois et al., 2006; Guthrie et al., 2013). Therefore, the indirect pathway and the external segment of the globus pallidus(GPe) are not included.

#### Architecture

Our model contains five main groups (see Fig. 3). Three of these groups are excitatory. These are the cortex (CTX), the thalamus (THL), and the subthalamic nucleus (STN). Two populations are inhibitory. They correspond to the sensory–motor territories of the striatum (STR) and the GPi. The model has been further tailored into three segregated loops (Alexander, DeLong, & Strick, 1986; Alexander & Crutcher, 1990; Alexander, Crutcher, & DeLong, 1991; Mink, 1996; Haber, 2003), namely the motor loop, the associative loop and the cognitive (or limbic) loop. The motor loop comprises the motor cortex (supplementary motor area (SMA), primary cortex (M1), premotor cortex (PMC), cingulate motor area (CMA)), the motor striatum (putamen), the motor STN, the motor GPi (motor territory of the pallidum and the substantia nigra) and the motor thalamus (ventrolateral thalamus (VLm and VLo)). The associative loop comprises the cognitive cortex (dorsolateral prefrontal cortex (DLPFC), the lateral orbitofrontal cortex (LOFC)) and the associative striatum (associative territory of the caudate). The cognitive loop comprises the cognitive cortex (anterior cingulate area (ACA), medial orbitofrontal cortex (MOFC)), the cognitive striatum (ventral caudate), the cognitive STN, the cognitive GPi (limbic territory of the pallidum and the substantia nigra and) the cognitive thalamus (ventral anterior thalamus (VApc, VAmc)).

**Figure 3.**
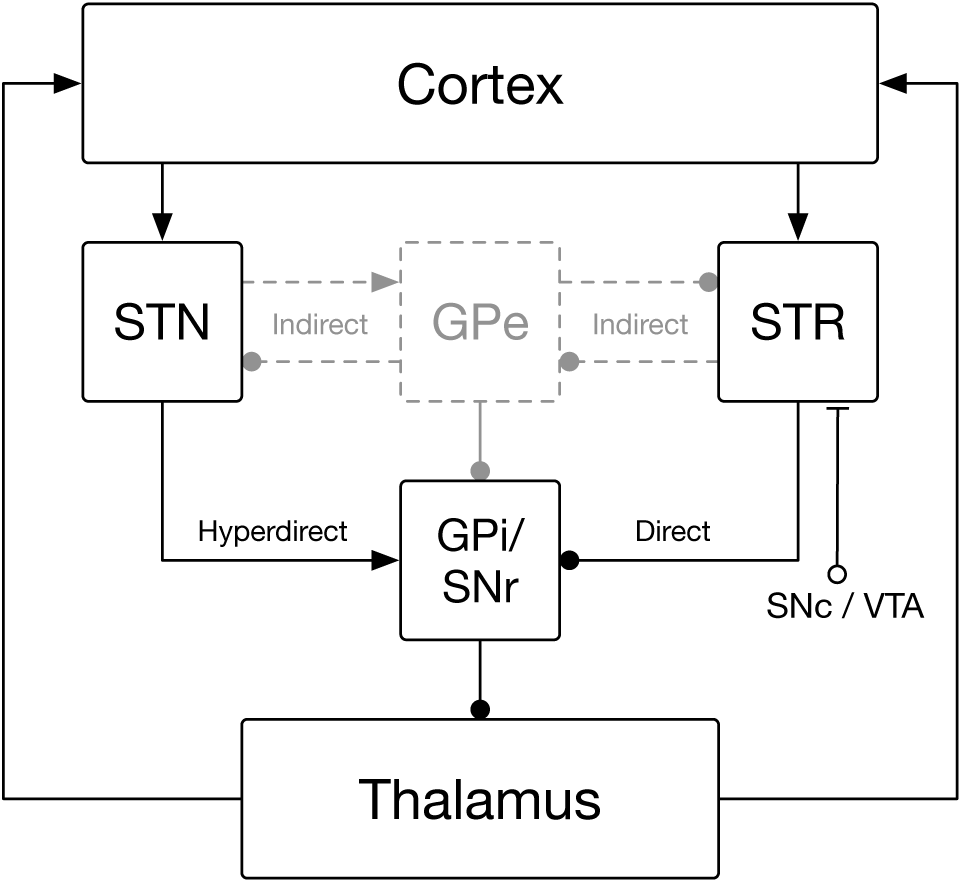
Architecture of the model. The architecture of the model is centered around the hyperdirect pathway (cortex → subthalamic nucleus → GPi/SNr → thalamus → cortex), the direct pathway (cortex → striatum → GPi/SNr → thalamus → cortex) and the cortex where lateral interactions take place (not represented on the figure). The external part of the globus pallidus, while not present in the model, is represented on the figure as a reminder of the actual connectivity in the BG. Similarly, the substantia nigra pars compacta is not explicitly represented in the model.

#### Populations

The model comprises 12 populations: 5 motor populations, 4 cognitive populations and 2 associative populations (Fig. 4). These populations comprises from 4 to 16 neural assemblies and possess each a specific geometry whose goal is to facilitate connectivity description. Each assembly is modeled using a neuronal rate model (Hopfield, 1984; Shriki, Hansel, & Sompolinsky, 2003) that give account of the spatial mean firing rate of the neurons composing the assembly. Each assembly is governed by the following equations:

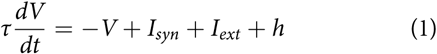

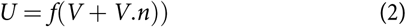

where *τ* is the assembly time constant (decay of the synaptic input), *V* is the firing rate of the assembly, *I*_*syn*_ is the synaptic input to the assembly, *I*_*ext*_ is the external input representing the sensory visual salience of the cue, *h* is the threshold of the assembly, *ƒ* is the transfer function and *n* is the (correlated, white) noise term. Each population possess its own set of parameters according to the group it belongs to (see Table 1). Transfer function for all population but the striatal population is a ramp function (*ƒ*(*x*) = *max*(*x*, 0)). The striatal population that is silent at rest (Sandstrom & Rebec, 2002), requires concerted coordinated input to cause firing (C. J. Wilson & Groves, 1981), and has a sigmoidal transfer function (nonlinear relationship between input current and membrane potential) due to both inward and outward potassium current rectification (Nisenbaum & Wilson, 1995). This is modeled by applying a sigmoidal transfer function to the activation of cortico-costriatal inputs in the form of the Boltzmann equation:

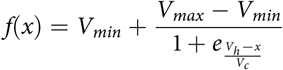

where *V*_*min*_ is the minimum activation, *V*_*max*_ the maximum activation, *V*_*h*_ the half-activation, and *V*_*c*_ the slope. This is similar to the use of the output threshold in the (Gurney, Prescott, & Redgrave, 2001) model and results in small or no activation to weak inputs with a rapid rise in activation to a plateau level for stronger inputs. The parameters used for this transfer function are shown in Table 2 and were selected to give a low striatal output with no cortical activation (1 spike/s), starting to rise with a cortical input of 10 sp/s and a striatal output of 20 spike/s at a cortical activation of 30 spike/s.

**Figure 4.**
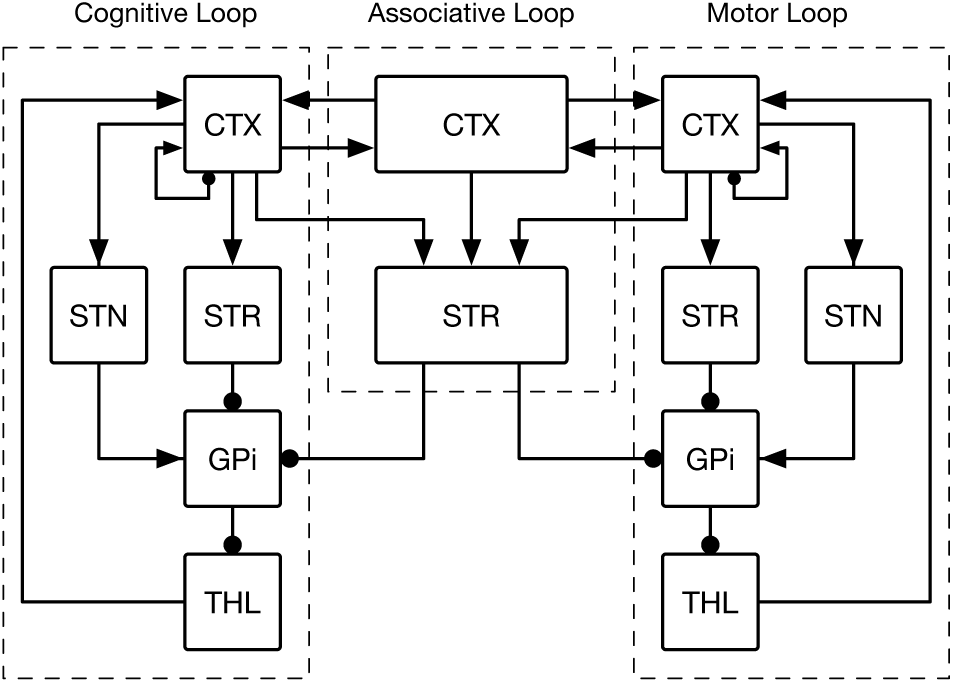
Segregated loops. The model is further detailed into three segregated circuits (cognitive, associative, motor). The cognitive and motor circuit each comprises a cortical, a striatal, a thalamic, a subthalamic, and a pallidal population while the associative loop only comprises a cortical and a striatal population. This latter interacts with the two other circuits via diffused connections to the pallidal regions and from all cortical populations. Arrows, excitatory connections. Dots, inhibitory connections.

**Table 1.** Population parameters

| Population | | Geometry | $\tau$ | Threshold | Noise |
| --- | --- | --- | --- | --- | --- |
| Cortex | associative | (4,4) | 10ms | -3 | 1.0% |
|  | cognitive | (4,1) | 10ms | -3 | 1.0% |
|  | motor | (1,4) | 10ms | -3 | 1.0% |
| Striatum | associative | (4,4) | 10ms | 0 | 0.1% |
|  | cognitive | (4,1) | 10ms | 0 | 0.1% |
|  | motor | (4,1) | 10ms | 0 | 0.1% |
| GPi | cognitive | (4,1) | 10ms | -10 | 3.0% |
|  | motor | (1,4) | 10ms | -10 | 3.0% |
| STN | cognitive | (4,1) | 10ms | -10 | 0.1% |
|  | motor | (1,4) | 10ms | -10 | 0.1% |
| Thalamus | cognitive | (4,1) | 10ms | -40 | 0.1% |
|  | motor | (1,4) | 10ms | -40 | 0.1% |

**Table 2.** Parameters for striatal sigmoid transfer function

| Name | Value |
| --- | --- |
| $V_{min}$ | 1 |
| $V_{max}$ | 20 |
| $V_h$ | 16 |
| $V_c$ | 3 |

#### Connectivity

Even though the model takes advantage of segregated loops, they cannot be entirely separated if we want the cognitive and the motor channel to interact. This is the reason why we incorporated a divergence in the corticostriatal connection followed by a re-convergence within the GPi (Graybiel, Aosaki, Flaherty, & Kimura, 1994; Parent et al., 2000) (see Fig. 5). Furthermore, we considered the somatotopic projection of the pyramidal cortical neurons to the striatum (Webster, 1961) as well as their arborization(Cowan & Wilson, 1994; C. J. Wilson, 1987; Parent et al., 2000; Parthasarathy, Schall, & Graybiel, 1992) resulting in specific localized areas of button formation (Kincaid, Zheng, & Wilson, 1998) and small cortical areas innervating the striatum in a discontinuous pattern with areas of denser innervation separated by areas of sparse innervation (Brown, Smith, & Goldbloom, 1998; Flaherty & Graybiel, 1991). We also cinsidered the large reduction in the number of neurons from cortex to striatum to GPi (Bar-Gad & Bergman, 2001; Oorschot, 1996). These findings combined lead to striatal areas that are mostly specific for input from one cortical area alongside areas where there is overlap between inputs from two or more cortical areas (Takada et al., 2001) and which are here referred to as the associative striatum.

**Figure 5.**
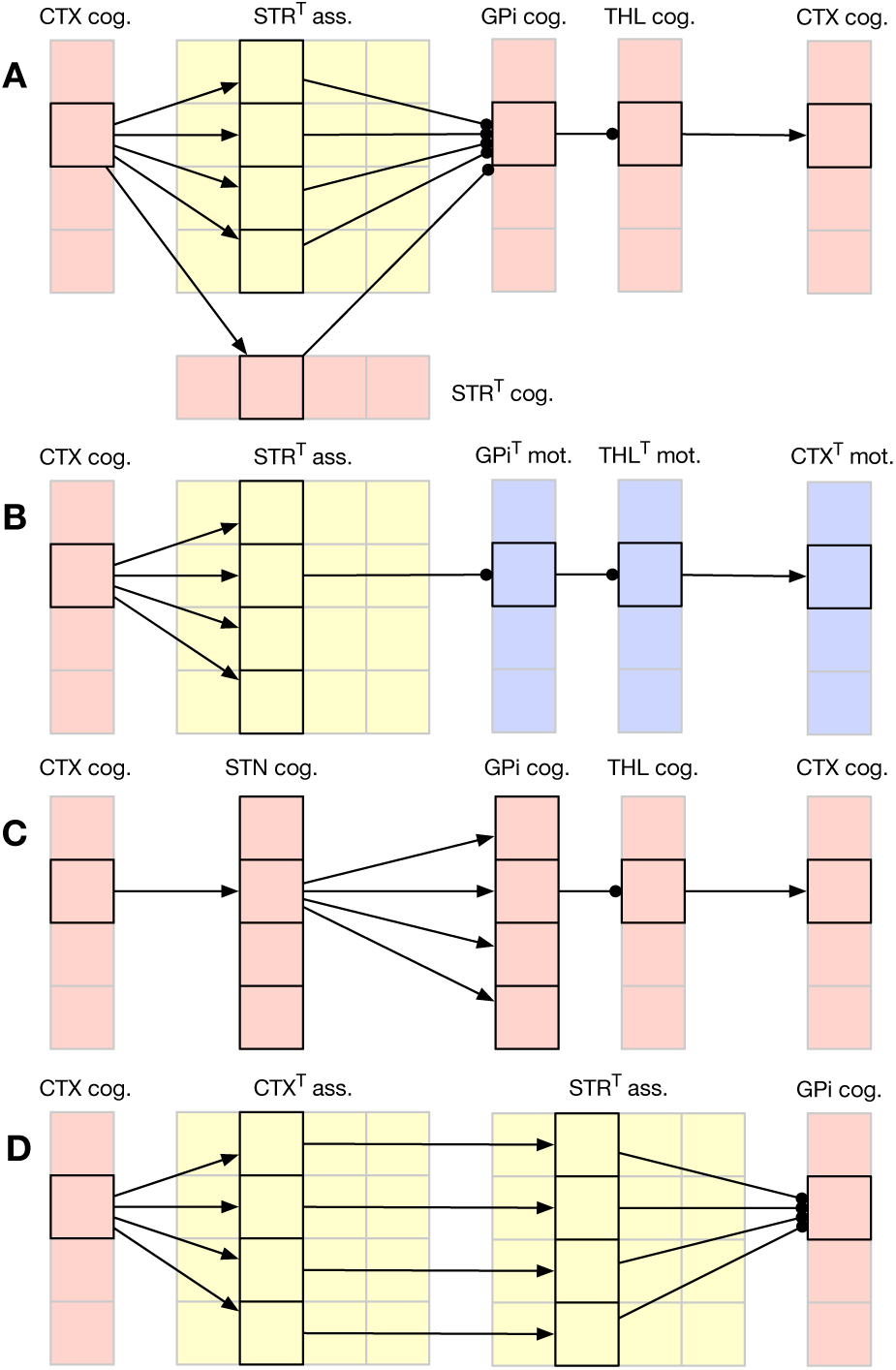
Partial connectivity in the cognitive and associative loops. For clarity, only one assembly has been considered. The motor loop is symmetric to the cognitive one. The “T” symbol on some name means the geometry of the group has been transposed (for readability). **A** The direct pathway from cognitive cortical assemblies diverge from cortex to associative and cognitive striatum. the pathway converges into cognitive GPi, send parallel projection to the thalamus and forms a closed loop with the original cognitive cortical assembly. **B** Thanks to the convergence of motor and cognitive pathways in association striatum, there is a cross-talking between the motor and cognitive loops. This allow a decision made in the cognitive loop to influence the decision in motor loops and vice-versa. **C** The hyperdirect pathway from cognitive cortical assembly diverges from STN to GPi, innervating all cognitive, but not motor, GPi regions and feeds back to all cognitive cortical assemblies. **D** The pathway from associative cortex and associative striatum is made of parallel localized projections.

The gain of the synaptic connection from population *A* (presy-naptic) to population *B* (postsynaptic) is denoted as *G*_*A*→*B*_, and the total synaptic input to population B is:

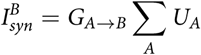

where *A* is the presynaptic assembly, *B* is the postsynaptic assembly, and *U*_*A*_ is the output of presynaptic assembly *A*. The gains for each pathway are shown in Table 3. Gains to the corresponding cognitive (motor) assembly are initially five times higher than to each receiving associative area. Re-convergence from cognitive (motor) and association areas of striatum to cognitive (motor) areas of GPi are evenly weighted.

**Table 3.** Connectivity gains and pattern between the different populations. For connectivity patterns, “*” means all. For example, (1,i) → (1,*) means one-to-all connectivity while (1,i) → (1,i) means one-to-one connectivity. Plastic pathways are indicated by a “•d” symbol.

| Pop. A | Pop. B | Pathway | Pattern | Gain |
| --- | --- | --- | --- | --- |
| Cortex | Striatum | cog. → cog. • | (i,1) → (i,1) | 1.0 |
|  |  | mot. → mot. | (i,1) → (i,1) | 1.0 |
|  |  | ass. → ass. | (i,j) → (i,j) | 1.0 |
|  |  | cog. → ass. | (i,1) → (i,*) | 0.2 |
|  |  | mot. → ass. | (1,i) → (*,i) | 0.2 |
|  | STN | cog. → cog. | (i,1) → (i,1) | 1.0 |
|  |  | mot. → mot. | (1,i) → (1,i) | 1.0 |
|  | Thalamus | cog. → cog. | (i,1) → (i,1) | 0.1 |
|  |  | mot. → mot. | (1,i) → (1,i) | 0.1 |
|  | Cortex | cog. → cog. | (i,1) → (*,1) | ±0.5 |
|  |  | mot. → mot. | (1,i) → (1,*) | ±0.5 |
|  |  | ass. → ass. | (i,j) → (*,*) | ±0.5 |
|  |  | ass. → mot. | (*,i) → (1,i) | 0.025 |
|  |  | ass. → cog. | (i,*) → (i,1) | 0.01 |
|  |  | cog. → ass. • | (i,1) → (i,*) | 0.025 |
|  |  | mot. → ass. | (1,i) → (*,i) | 0.01 |
| Striatum | GPi | cog. → cog. | (i,1) → (i,1) | -2.0 |
|  |  | mot. → mot. | (1,i) → (1,i) | -2.0 |
|  |  | ass. → cog. | (i,*) → (i,1) | -2.0 |
|  |  | ass. → mot. | (*,i) → (1,i) | -2.0 |
| STN | GPi | cog. → cog. | (i,1) → (i,1) | 1.0 |
|  |  | mot. → mot. | (1,i) → (1,i) | 1.0 |
| GPi | Thalamus | cog. → cog. | (i,1) → (i,1) | -1.0 |
|  |  | mot. → mot. | (1,i) → (1,i) | -1.0 |
| Thalamus | Cortex | cog. → cog. | (i,1) → (i,1) | 1.0 |
|  |  | mot. → mot. | (1,i) → (1,i) | 1.0 |

#### Task encoding

At the trial start, assemblies in the cognitive cortex encoding the two cues *C*_1_ and *C*_2_ receive an external current (7Hz) and assemblies in the motor cortex encoding the two positions *M*_1_ and *M*_2_ receive similarly an external current (7Hz). These activities are not sufficient to disambiguate between the situation (*C*_1_/*M*_1_, *C*_2_/*M*_2_) and the situation (*C*_1_/*M*_2_, *C*_2_/*M*_1_). This is the reason why the associative cortex encoding one of these two situations receives an external current (7Hz), (*C*_1_/*M*_1_, *C*_2_/*M*_2_) in the present case (see Fig. 6. The decision of the model is decoded from the activity in the motor cortex *only*, i.e. independently of the activity in the cognitive cortex. If the model chooses a given cue but produces the wrong motor command, the cognitive choice will not be taken into account and the final choice will be decoded from the motor command that may lead to an irrelevant choice.

**Figure 6.**
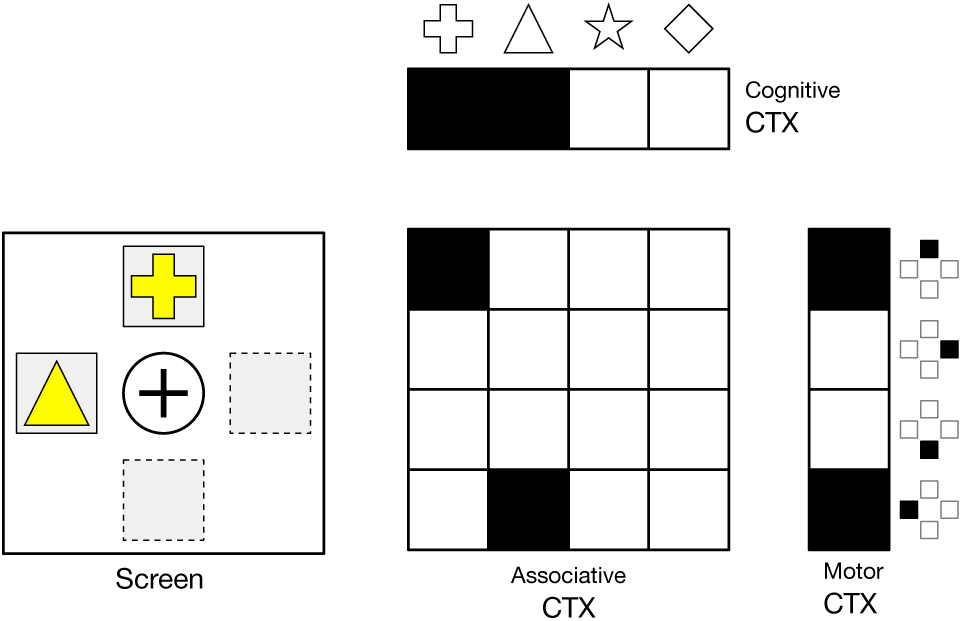
Task encoding. Assemblies in the cognitive cortex encoding the two cues *C*_1_ and *C*_2_ receive an external current and assemblies in the motor cortex encoding the two positions *M*_1_ and *M*_2_ receive similarly an external current. These activities are not suffcient to disambiguate between the situation (*C*_1_/*M*_1_, *C*_2_/*M*_2_) and the situation (*C*_1_/*M*_2_, *C*_2_/*M*_1_). This is the reason why the associative cortex encoding one of these two situations receives an external current, (*C*_1_/*M*_1_, *C*_2_/*M*_2_) in the present case.

#### Dynamic

There exist two different competition mechanisms inside the model. One is conveyed through the direct and hyperdirect pathways, the other is conveyed inside the cortex through short-range excitation and long range inhibition. The former has been fully described and analyzed in Leblois et al., 2006 while the latter been extensively studied in a number of experimental and theoretical papers (von der Malsburg, 1973; H. R. Wilson & Cowan, 1972, 1973; Amari, 1977; Callaway, 1998; Taylor, 1999). Each of these two competition mechanisms can lead to a decision as illustrated on Fig. 7 that shows the dynamic of the motor loop for all the population in three conditions. In the absence of the cortical interactions (gain of cortical lateral connections has been set to 0), the direct and hyperdirect pathway are able to promote a competition that result in the selection of one of the two assemblies in each group. In the absence of GPi output (connection has been cut), the cortical lateral connections are able to support a competition that result in the selection of o the two assemblies, even though such decision is generally slower than the basal one. The result of the dual competition is a faster selection of one of the two assemblies prior to learning, when there is no possibility for the two competition to be non congruent (one competition tends to select move A while the others tend to select move B). We’ll see in the results section that if the result of the two competitions is non-congruent, the decision is slower.

**Figure 7.**
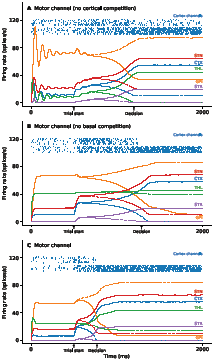
Activity in the different populations during a single trial of action selection before learning. The model is started at time t=0ms and allowed to settle to a steady state before the presentation of the cues at t=500ms. Solid lines represents activity related to the selected population, dashed lines represent activity related to the non selected population. Decision threshold has been set to 40 spikes/s between the two cortical populations and is indicated on the x axis. Raster plots are related to the cortical populations and has been generated from the firing rate of 10 neurons. **A** Activity in the motor populations in the absence of lateral competition in the cortical populations. The damped oscillations during the settling phase are characteristic of the delayed feedback from the subthalamic nucleus (excitation) and the striatum (inhibitory) through the globus pallidus and the thalamus. **B** Activity in the motor populations in the absence of the feedback from the basal ganglia (GPi) to the cortical populations via the thalamus. Decision threshold is reached thanks to the direct lateral competition in both cognitive and motor cortical channels. There is no damped oscillation since there is no delay between the cortical populations and the decision times are slower than in the previous case. **C** Activity in the motor populations in the full model with a dual competition, one cortical, one basal. When congruent (cortical and basal decision are the same), decision time for both the motor and cortical channels are faster than in the absence of one of the competition loop.

#### Learning

Learning has been restricted to the cognitive channel on the cortico-striatal synapse (between the cortex cognitive and the striatum cognitive) and the cortico-cortical synapse (between the cortex cognitive and the cortex associative). There is most probably learning in other structures and pathways, but the aim here is to show that the proposed restriction is sufficient to produce the behavior under consideration. All synaptic weights are initialized to 0.5 (SD 0.005) that are used as as a multiplier to the pathway gain to keep the factors of gain and weight separately observable. All weights are bound between *W*_*min*_ and *W*_*max*_ (see Table 4) such that for any change Δ*W*(*t*), weight *W*(*t*) is updated according to the equation:

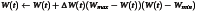

**Table 4.** Learning parameters

| Name | Value |
| --- | --- |
| $W_{min}$ | 0.25 |
| $W_{max}$ | 0.75 |
| $LTP_{RL}$ | 0.050 |
| $LTD_{RL}$ | 0.030 |
| $LTP_{HL}$ | 0.005 |
| $\alpha$ | 0.025 |

##### Reinforcement learning

At the level of corticostriatal synapses, phasic changes in dopamine concentration have been shown to be necessary for the production of long-term potentiation (LTP) (Kerr & Wickens, 2001; Reynolds, Hyland, & Wickens, 2001; Surmeier, Ding, Day, Wang, & Shen, 2007; Pawlak & Kerr, 2008). After each trial, once reward has been received (0 or 1), the corticostriatal weights are updated according to:

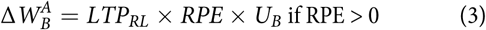

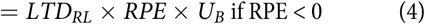

where 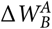 is the change in the weight of the corticostriatal synapse from cortical assembly A to striatal assembly B, *RPE* is the reward prediction error, the amount by which the actual reward delivered differs from the expected reward, *U*_*B*_ is the activation of the striatal assembly, and *μ* is the actor learning rate. Generation of LTP and long-term depression (LTD) in striatal MSNs has been found to be asymmetric (Pawlak & Kerr, 2008). Therefore, in the model, the actor learning rate is different for LTP and LTD. The RPE is calculated using a simple critic learning algorithm:

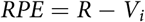

where R, the reward, is 0 or 1, depending on whether a reward was given or not on that trial. Whether a reward was given was based on the reward probability of the selected cue (which is most of the time the one associated with the direction chosen). i is the number of the cue chosen, and *V*_*i*_ is the value of cue i. The value of the chosen cue is then updated using the PE:

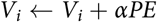

##### Hebbian learning

At the level of cortico-cortical synapse, only the co-activation of two assemblies is necessary for the production of long-term potentiation (Bear & Malenka, 1994; Caporale & Dan, 2008; Feldman, 2009; Hiratani & Fukai, 2016). After each trial, once a move has been initiated, the cortico-cortical weights are updated according to:

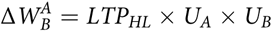

where 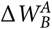 is the change in the weight of the cortico-cortical synapse from cognitive cortical assembly A to associative cortical assembly B. This learning rule is thus independent of reward.

### Experimental setup

*With kind permission from the authors (Piron et al., 2016), we reproduce here the details of the experimental setup as well as the surgical procedure since the two same monkeys were used for these new experiments*.

Experimental data were obtained from 2 female macaque monkeys (Macaca mulata). Experiments were performed during the daytime. Monkeys were living under a 12h/12h diurnal rhythm. Although food access was available ad libitum, the primates were kept under water restriction to increase their motivation to work. A veterinary skilled in healthcare and maintenance in nonhuman primates supervised all aspects of animal care. Experimental procedures were performed in accordance with the Council Directive of 20 October 2010 (2010/63/ UE) of the European Community. This project was approved by the French Ethic Comity for Animal Experimentation (50120111-A).

#### Surgical Procedure

Cannula guides were implanted into the left and right GPi in both animals under general anesthesia. Implantation was performed inside a stereotaxic frame guided by ventriculography and single-unit electrophysiological recordings. A ventriculographic cannula was introduced into the anterior horn of the lateral ventricle and a contrast medium was injected. Corrections in the position of the GPi were performed according to the line between the anterior commissure (AC) and the posterior commissure (PC) line. The theoretical target was AP: 23.0mm, L: 7.0 mm, P: 21.2 mm.27 A linear 16-channel multielectrode array was lowered vertically into the brain. Extracellular single-unit activity was recorded from 0mm to 24 mm relative to the AC–PC line with a wireless recording system. Penetration of the electrode array into the GPi was characterized by an increase in the background activity with the appearance of active neurons with a tonic firing rate (around the AC–PC line). The exit of the electrode tips from the GPi was characterized by the absence of spike (around 3-4 mm below the AC–PC line). When a clear GPi signal from at least 3 contacts had been obtained, control radiography of the position of the recording electrode was performed and compared to the expected position of the target according to the ventriculography. If the deviation from the expected target was less than 1mm, the electrode was removed and a cannula guide was inserted with a spare cannula inside so that the tip of the cannula was superimposed on the location of the electrode array in the control radiography. Once the cannula guide was satisfactorily placed, it was fixed to the skull with dental cement.

#### Bilateral Inactivation of the GPi

Micro-injections were delivered bilaterally 15 minutes before a session. For both animals injections of the *GABA*_*A*_ agonist muscimol hydrobromide (Sigma) or saline (NaCl 9‰) were randomly assigned each day. Muscimol was delivered at a concentration of 1 *μ*g/*μ*l (dissolved in a NaCl vehicle). Injections (1*μ*l in each side) were performed at a constant flow rate of 0.2 *μ*l/min using a micro-injection system. Injections were made through a 30-gauge cannulae inserted into the 2 guide cannulae targeting left and right GPi. Cannulas were connected to a 25 *μ*l Hamilton syringe by polyethylene cannula tubing.

### Data Analysis

Theoretical and experimental data were analyzed using Kruskal-Wallis rank sum test between the three conditions (saline (C0), muscimol (C1) or saline following muscimol (C2)) for the 6 samples (12×10 first trials of C0 (control), 12×10 last trials of C0 (control), 12×10 first trials of C1 (GPi Off/muscimol); 12×10 last trials of C1(GPi OFF/muscimol); 12×10 first trails of C2(GPi On/saline); 12×10 last trials of C2(GPi On/saline)) with posthoc pairwise comparisons using Dunn’s-test for multiple comparisons of independent samples. P-values have been adjusted according to the false discovery rate (FDR) procedure of Benjamini-Hochberg. Results were obtained from raw data using the PMCMR R package (Pohlert, 2014). Significance level was set at *P* < 0.01.

Experimental raw data is available from (Kase & Boraud, 2017) under a CC0 license, Theoretical raw data and code are available from (Rougier & Topalidou, 2017) under a CC0 license (data) and BSD license (code).

## Results

Our model predicts that the valuation of options and the behavioural outcome are two separate (but entangled) processes. This means that if we block the output of the basal ganglia before learning, reinforcement learning still occurs at the striatal level under dopaminergic control and this should induce covert learning of stimuli value even though the behavioral choice would appear as random.

### Protocol

The protocol has been consequently split over two consecutive conditions (C1 & C2) using the same set of stimuli and a dissociated control (C0) using a different set of stimuli (using same probabilities as for C1 & C2).

> C0 60 trials, GPi On (model), saline injection (primates), stimulus set 1 (*A*_1_, *B*_1_) with *P*_*A*_1__ = 0.75, *P*_*B*_1__ = 0.25
>
> C1 60 trials, GPi Off (model), muscimol injection (primates), stimulus set 2 (*A*_2_, *B*_2_) with *P*_*A*_2__ = 0.75, *P*_*B*_2__ = 0.25
>
> C2 60 trials, GPi On (model), saline injection (primates), stimulus set 2 (*A*_2_, *B*_2_) with *P*_*A*_2__ = 0.75, *P*_*B*_2__ = 0.25

### Computational results

We tested our hypothesis on the model using 12 different sessions (corresponding to 12 different initializations of the model). On day 1, we suppressed the GPi output by cutting the connections between the GPi and the thalamus. When the GPi output has been suppressed on day 1, the performance is random at the beginning as shown by the average probability of choosing the best option (expressed in mean±SD) in the first 10 trials (0.408±0.161) and remain so until the end of the session (0.525 ±0.164). Statistical analysis revealed no significant difference between the 10 first and the 10 last trials. On day 2, we re-established connections between the GPi and the thalamus and the model has to perform the exact same task as for day1 using the same set of stimuli. Results shows a significant change in behavior: the model starts with an above-chance performance on the first 10 trials (0.717±0.241) and this change is significant (see Table 5 and Fig. 8) as compared to the start of C1, as compared to the end of C1 and as compared to the start of C0, confirming our hypothesis that the BG have previously learned the value of stimuli even though they were unable to alter behavior.

**Table 5.** Theoretical results statistical analysis. Kruskal-Wallis rank sum test between the three conditions (saline (C0), muscimol (C1) or saline following muscimol (C2)) with posthoc pairwise comparisons using Dunn’s-test for multiple comparisons of independent samples.

| H0 | statistic (H) | p value |
| --- | --- | --- |
| C0 start = C2 start | 2.965 | 0.0051 |
| C1 start = C2 start | 4.986 | 1.8e-6 |
| C1 end = C2 start | 3.099 | 0.0036 |

**Figure 8.**
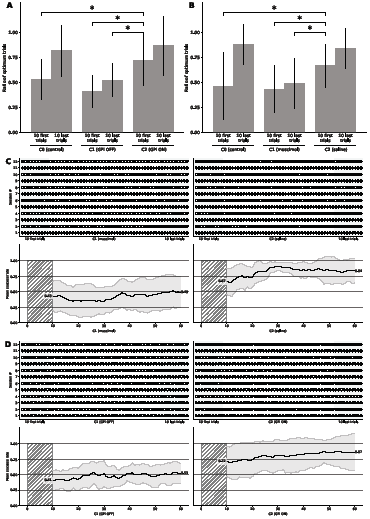
Theoretical and experimental results. Histograms show the mean performance at the start and the end of a session in day 1 and day 2 conditions for both the model (**A**) and the monkeys (**B**). At the start of day 2, the performance for both the model and the monkeys is significantly higher compared to the start and end of day 1, suggesting some covert learning occurred during day 1 even though performances are random during day 1. **C** Individual trials (n=2x60) for all the sessions (n=12) for the primates. **D** Individual trials (n=2x60) for all the sessions (n=12) for the model. A black dot means a successful trial (the best stimulus has been chosen), an outlined white dot means a failed trial (the best stimulus has not been chosen). Measure of success is independent of the actual reward received after having chosen one of the two stimuli. The bottom part of each panel shows the mean success rate over a sliding window of ten consecutive trials and averaged across all the sessions. The thick black line is the actual mean and the gray-shaded area represents the standard deviation (STD) over sessions.

### Experimental results

We tested the prediction on two female macaque monkeys which have been implanted with two cannula guides into the left and right GPi (see Materials and Methods section for details). In order to inhibit the GPi, we injected bilaterally a GABA agonist (muscimol, 1*μ*g) 15 minutes before working session on day 1 (C1). The two monkeys were trained for 7 and 5 sessions respectively, each session using the same set of stimuli. Results on day 1 shows that animals were unable to choose the best stimulus in such condition from the start (0.433 ±0.236) to the end (0.492±0.250) of the session. Statistical analysis revealed no significant difference between the 10 first and the 10 last trials on day 1. On day 2 (C2), we injected bilaterally a saline solution 15 minutes before working session and animals had to perform the exact same protocol as for day 1. Results shows a significant change in behavior (see Table 6 and Fig. 8): animals start with an above-chance performance on the first 10 trials (P=0.667 ±0.213), as compared to the start of C1, as compared to the end of C1 and as compared to the start of C0, confirming our hypothesis that the BG has previously learned the value of stimuli.

**Table 6.**
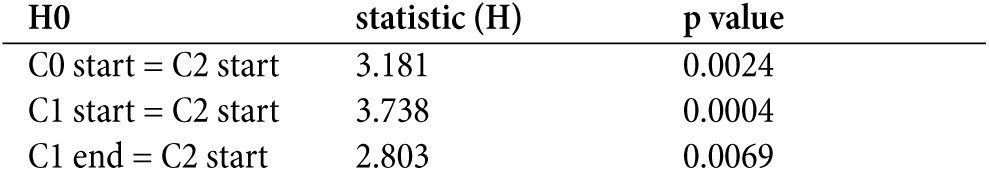
Experimental results statistical analysis. Kruskal-Wallis rank sum test between the three conditions (saline (C0), muscimol (C1) or saline following muscimol (C2)) with posthoc pairwise comparisons using Dunn’s-test for multiple comparisons of independent samples.

| H0 | statistic (H) | p value |
| --- | --- | --- |
| C0 start = C2 start | 3.181 | 0.0024 |
| C1 start = C2 start | 3.738 | 0.0004 |
| C1 end = C2 start | 2.803 | 0.0069 |

## Discussion

### Covert learning in the BG

These results reinforce the classical idea that the basal ganglia architecture is based on an actor critic architecture where the dopamine serves as a reinforcement signal. However, the proposed model goes beyond this classical hypothesis and proposes a more general view on the role of the BG in behaviour and the entanglement with the cortex. Our results, both theoretical and experimental, suggest that the critic part of the BG extends its role beyond the basal ganglia and makes it *de facto* a central component in behavior that can evaluate any action, independently of their origin. This hypothesis is very congruent with the results introduced in Charlesworth, Warren, and Brainard (2012) where authors show that the anterior forebrain pathway in Bengalese finches contributes to skill learning even when it is blocked and does not participate in the behavioural performance. This is also quite compatible with (Ashby, Turner, & Horvitz, 2010; Hélie, Ell, & Ashby, 2015) who propose that the BG is a general purpose trainer for cortico-cortical connections. Here, we introduced a precise computational model using both reinforcement and Hebbian learning, supported by experimental data, that explains precisely how this general purpose trainer can be biologically implemented.

This can be simply understood by scrutinizing a session in control and lesion condition (see Fig. 9). In control condition, the model learns to select the best cue thanks to the BG. Because it learns what is the best stimulus, this induces a preferential selection of the best stimulus in order to obtain a higher probability of reward. If the process is repeated over many trials, this leads implicitly to an over-representation of the more valuable stimuli at the cortical level and since cortex learns with Hebbian learning, it is implicitly learned. Said differently, the value of the best stimulus has been *converted* to the temporal domain. In lesion condition, the selection is random and each stimulus is roughly selected with equal probability and this allows the BG to evaluate the value of the two stimuli even more precisely. We believe this is the same for the monkeys even though we do not have access to internal value and weights. However, we can see on Fig. 10 that the estimated value of stimuli (computed as the probability of reward) reflects the highest value for the best stimulus. Similarly, the number of time a given stimulus has been selected is correlated with its actual value even if it is not significant.

**Figure 9.**
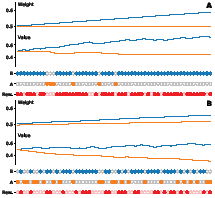
Model performance during a single session. Filled dots indicate the chosen cue between A and B. Filled red dots indicate if a reward has been received following the choice. Reward probability is 0.75 for cue A and 0.25 for cue B. **A** Intact model (C0). The BG output drives the decision and evaluates the value of cue A and cue B with a strong bias in favor of A because this cue is chosen more frequently. In the meantime, the Hebbian weight relative to this cue is strongly increased while the weight relative to the other cue doesn’t change significantly. **B** Lesioned model (C1). The BG output has been suppressed and decisions are random. Hebbian weights for cue A and cue B are both increased up to similar values at the end of the session. In the meantime, the value of cue A and cue B are evaluated within the BG and the random sampling of cue A and cue B leads to an actual better sampling of value A and B. This is clearly indicated by the estimated value of B that is very close the theoretical value (0.25).

**Figure 10.**
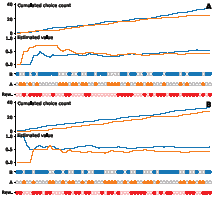
Monkey performance during a single session. Filled dots indicate the chosen cue between A and B. Filled red dots indicate if a reward has been received following the choice. Reward probability is 0.75 for cue A and 0.25 for cue B. **A** In saline condition (C0), the monkey is able to slowly choose for the best cue with a slight preferences for A at the end of the 60 trials. Estimation of the perceived value of the two cues shows the actual value of A is greater than the value of B at the end of the session **B** In muscimol condition (C1), the monkey choose cues randomly as indicated by the overall count of choices A and B. Estimation of the perceived value of the two cues (dashed lines) reveals a greater estimation of the value of A compared to the value of B.

### From action-outcome to stimulus-response

These new results, together with our previous results (Piron et al., 2016) shed a new light on a plausible neural mechanism responsible for the gradual mix between an action-outcome behavior and a stimulus-response one. The novelty in our hypothesis is that there is no transfer *per se*. There is instead a joint combination of the two systems that act and learn together and we tend to disagree with the hypothesis of a hierarchical system (Dezfouli & Balleine, 2013). In our case, the final behavioral decision results from a subtle balance between the two decisions. When a new task needs to be solved, the basal ganglia initially drives the decision because it has initially a faster dynamic. In the meantime, the cortex takes advantage of this driving and gradually learns the decision independently of the reward. We’ve shown how this could be the case for monkeys, even though we lack experimental evidence that the decision in muscimol condition is actually driven by the cortex. The actual combination of the two systems might be more complex than a simple weighted linear combination and this make the study even more difficult to carry on. What we see at the experimental level might the projection of a more complex phenomenon. Persisting in a devaluated task does not mean the system is *frozen* but the time to come back from a stimulus-response oriented behavior might be simply much longer than the time to initially acquire the behavior.

Finally, our results suggest a behavioral decision results from both the cooperation (acquisition) and competition (expression) of two distinct but entangled systems.

## Abbreviations

**A-O** Action – Outcome

**AC** Anterior Commissure

**CMA** Cingulate motor area

**CTX** Cortex

**DLPFC** Dorsolateral prefrontal cortex

**DLS** Dorsolateral striatum

**DMS** Dorsomedial striatum

**FEF** Frontal eye fields

**GPi** Internal part of the globus pallidus

**GPe** External part of the globus pallidus

**LTP** Long-term potentiation

**LTD** Long-term depression

**LOFC** Lateral orbitofrontal cortex

**M1** Primary motor cortex

**MOFC** Medial orbitofrontal cortex

**OFC** Orbitofrontal cortex

**PC** Posterior commissure

**PFC** Prefrontal cortex

**PMC** Premotor cortex

**RPE** Reward prediction error

**SMA** Supplementary motor area

**SNc** Substantia nigra pars compacta

**SNr** Substantia nigra pars reticulata

**STN** Subthalamic nucleus

**STR** Striatum

**S-R** Stimulus – Response

**THL** Thalamus

**VLm** Ventrolateral thalamus, pars medialis

**VLo** Ventrolateral thalamus, pars oralis

**VApc** Ventral anterior thalamus, pars parvocellularis

**VAmc** Ventral anterior thalamus, pars magnocellularis

## Footnotes

1 For example on Fig. 1, if we consider a uniform cue sampling policy for 6*n trials, the mean expected reward for a perfect player with full knowledge is 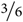 × 1 + 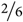 × ⅔ + ⅙ × ⅓ = 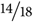 ≈ 0.777…

